# Daily changes in neuronal activities of the dorsal motor nucleus of the vagus under standard and short-term high fat diet – implications for circadian modulation of parasympathetic outflow

**DOI:** 10.1101/2021.03.02.433571

**Authors:** Lukasz Chrobok, Jasmin D Klich, Jagoda S Jeczmien-Lazur, Kamil Pradel, Katarzyna Palus-Chramiec, Anna M Sanetra, Hugh D Piggins, Marian H Lewandowski

## Abstract

The suprachiasmatic nuclei (SCN) of the hypothalamus functions as the brain’s primary circadian clock, but circadian clock genes are also rhythmically expressed in several extra-SCN brain sites where they can exert local temporal control over physiology and behaviour. Recently, we found that the hindbrain dorsal vagal complex possesses strong daily timekeeping capabilities, with the area postrema and nucleus of the solitary tract exhibiting the most robust clock properties. The possibility that the executory part of this complex – the dorsal motor nucleus of the vagus (DMV), also exhibits daily changes has not been extensively studied. The DMV is the source of vagal efferent motoneurons largely responsible for the regulation of gastric motility and emptying and consequently influence meal size and energy homeostasis. We used a combination of multi-channel electrophysiology and patch clamp recordings to gain insight into possible daily variation in these DMV cells and how this is influenced by diet. We found that DMV neurons increase their spontaneous activity, excitability and responsiveness to metabolic neuromodulators at late day which was paralleled with an enhanced synaptic input to these neurons. A high-fat diet typically damps circadian rhythms, but we found that short-term exposure to a high-fat diet paradoxically amplified daily variation of DMV neuronal activity, while blunting their responsiveness to metabolic neuromodulators. In summary, we show for the first time that neural activity at a source of vagal efferents varies with time of day and that this temporal variation is modulated by diet. These findings have clear implications for our understanding of the daily control of parasympathetic outflow.

## 1. INTRODUCTION

Circadian rhythms have evolved as a mechanism by which life forms on our planet can anticipate and adapt to the recurrent 24 h variation in environmental conditions that arises from the Earth’s rotation on its axis. In mammals, internal timekeeping is observed at all levels of physiology – from daily variation in gene expression to complex behaviours such as the timing of feeding, drinking, or the onset of sleep. These rhythmic changes are governed by the circadian timing system with the suprachiasmatic nuclei of the hypothalamus (SCN) schematised as the master clock (Takahashi, 2017; Hastings *et al*., 2019). However, accumulating evidence challenges this uni-clock view, with several extra-SCN brain sites exhibiting rhythmic clock gene expression to potentially act as independent circadian clocks (Guilding & Piggins, 2007; Flanagan *et al*., 2020).

Recently, we reported unusually robust circadian timekeeping properties of the mouse dorsal vagal complex (DVC), a multi-component hindbrain centre which regulates a plethora of homeostatic processes including osmoregulation, satiety, parasympathetic tone and cardio-vascular function (Grill & Hayes, 2012; Chrobok *et al*., 2020). This complexity of DVC function is supported by its three discrete, yet closely linked neuronal structures: (1) the area postrema (AP), a sensory circumventricular organ, (2) the nucleus of the solitary tract (NTS), a powerful hub for visceral and cardiovascular cues, and (3) the dorsal motor nucleus of the vagus (DMV). The DMV is the executive part of the DVC and is the origin of preganglionic vagal motoneurons which drive both inhibitory and excitatory control over gastro-intestinal smooth muscle (Grill & Hayes, 2009, 2012; Browning & Travagli, 2014; Constantinescu, 2016). Neuronal activity in the AP/NTS peaks at late day/early night, coinciding with elevated expression of the clock gene, *Per2*, in these structures. In *ex vivo* investigation, *Per2* expression in cultured brain slices was robust and sustained for up to a week in the AP and NTS, whereas in the DMV, *Per2* expression was only transient (lasting one day). Further, while firing activity of AP and NTS neurons varies over 24 h *ex vivo*, such daily variation could not be assessed in the mouse DMV owing to difficulty in distinguishing DMV from NTS recording sites in those multi-electrode array recordings (Chrobok *et al*., 2020).

Current evidence supports the view of a functional organisation of the DVC by which its distinct nuclei play different roles in gastro-intestinal signal processing (Grijalva & Novin, 1990; Rogers *et al*., 1996; Grill & Hayes, 2012). Excitatory sensory vagal afferents terminate predominately in the NTS, which integrates gastric cues with central metabolic and homeostatic information (Kaelberer *et al*., 2018; Han *et al*., 2018). The principal target of this processed signal is the subjacent DMV, which receives mostly tonic inhibitory (but also some excitatory) signals from the NTS (Davis *et al*., 2004). Modulation of these NTS to DMV synapses exerts profound effects upon visceral function (Derbenev *et al*., 2004; Cruz *et al*., 2007). Thus, the NTS controls vagal outflow from the DMV through altering its synaptic input to this executory part of the DVC. Neuronal activity of DMV neurons is also potently modulated by metabolic signals that exhibit both periprandial and circadian rhythmicity. Appetite promoting factors include the hypothalamic orexin neuropeptides whose levels are increased preprandially and whose application to the DMV stimulates gastric motor function (Krowicki *et al*., 2002). In contrast, the anorexigenic glucagon-like peptide 1 (GLP-1) which is expressed in intestinal L-cells as well as NTS neurons, is released postprandially and acts via the DMV to induce gastroinhibition (Holmes *et al*., 2009). Both orexins and GLP-1 excite preganglionic DMV neurons, with their opposing effects on gastric motility believed to be segregated at the postganglionic level (Cruz *et al*., 2007).

Dietary constituents and dietary habits can modulate central nervous system function, including circadian timekeeping (Challet, 2013, 2015, 2019). The dysregulation of intrinsic circadian rhythmicity can result in metabolic syndrome and obesity, and conversely, consumption of calorie-dense high-fat diet alters endogenous rhythmic daily processes (Baron & Reid, 2014, 2015; Namvar *et al*., 2016; Engin, 2017). Typically such effects of diet are attributed to actions on the hypothalamus, but high-fat diet can affect the NTS and blunt or eliminate circadian rhythmicity in its clock gene expression (Kaneko *et al*., 2009; Zhang *et al*., 2020). In a companion study, we also show that even short-term consumption of a high-fat diet alters neuronal activities of the rat NTS (Chrobok *et al*., 2021*b*). Further, multiple studies report adverse effects of perinatal and adult exposure to high-fat diet on DMV neuronal activity, including disturbances in glutamatergic and GABAergic signalling and its sensitivity to neuropeptides (Browning *et al*., 2013; Bhagat *et al*., 2015; Clyburn *et al*., 2018, 2019). However, whether the DMV expresses daily rhythms and if so, whether such rhythms are influenced by diet is unknown.

To address this gap in current knowledge, we investigated daily variation in rat DMV neuronal activities, including firing rate, synaptic input and other fundamental membrane properties. We also assessed potential day-night difference in the responsiveness of DMV neurons to metabolically relevant neuropeptides. Additionally, we studied effects of short-term exposure to high-fat diet on these parameters. We report new evidence that there is a late day upregulation of DMV neural activities that manifests in increased synaptic input (predominately excitatory), enhanced excitability and spontaneous firing rates and in increased responsiveness to periprandial neuropeptides. Surprisingly, we found short-term high-fat diet to boost daily change in spontaneous activity and excitability of DMV neurons, while blunting their responsiveness to metabolic cues. Collectively, this study presents the first evidence that DMV neurons increase their activity and responsiveness around the day-night transition. These findings raise the possibility that daily changes in the DMV contribute to circadian variation in parasympathetic outflow.

## 2. MATERIALS AND METHODS

### 2.1. Ethical approval

All procedures were approved by the Local (Krakow) Ethical Commission and performed in accordance with the European Community Council Directive of 24 November 1986 (86/0609/ EEC) and the Polish Animal Welfare Act of 23 May 2012 (82/2012). Every effort was made to minimise the number of animals used in the study and their suffering.

### 2.2. Animals

This study was performed on 4-8 week old male Sprague Dawley rats housed at the Institute of Zoology and Biomedical Research animal facility at the Jagiellonian University in Krakow under constant environmental conditions (temperature: 23°C, humidity: ∼60%). Rats were kept under standard (12h/12h) light/dark cycle with water and food provided *ad libitum*. In these conditions, Zeitgeber time 12 (ZT12) represents lights-off and ZT0, lights-on.

### 2.3. Diet

Two different dietary conditions were introduced at postnatal day 28, when animals were weaned. Control diet (CD, ∼ 3,514 kcal/kg, energy from: 10% fat, 24% protein, 66% carbohydrates; Altromin, Germany) or high fat diet (HFD, ∼ 5,389 kcal/kg, energy from: 70% fat, 16% protein, 14% carbohydrates; Altromin) was given *ad libitum* for 2-3 weeks prior electrophysiological procedures, or for 4 weeks before the cull for immunohistochemistry.

### 2.4. Electrophysiology

#### 2.4.1. Tissue preparation

Electrophysiological experiments were conducted on brainstem coronal slices containing the DMV at the level of AP (anteroposterior -14.3 to -13.7, mediolateral -1.6 to +1.6, dorsoventral -7.6 to -8 mm from Bregma; Paxinos & Watson, 2007). Rats were culled at two distinct time points for patch clamp experiments (ZT3 and 15) and at four for multi-electrode array (MEA) recordings (ZT3, 9, 15 and 21). Animals were deeply anaesthetised with isoflurane inhalation (2 ml/kg of body weight) and decapitated. Then, brains were immediately removed from the skull and transferred to ice-cold preparation artificial cerebrospinal fluid (ACSF) composed of (in mM): 25 NaHCO_3_, 3 KCl, 1.2 Na_2_HPO_4_, 2 CaCl_2_, 10 MgCl_2_, 10 glucose, 125 sucrose with addition of pH indicator, Phenol Red 0.01 mg/l, osmolality ∼290 mOsmol/kg. ACSF was continuously bubbled with carbogen (95% O_2_, 5% CO_2_). Block of tissue containing the brainstem and cerebellum was then mounted on a cold plate of the vibroslicer (Leica VT1000S, Heidelberg, Germany) and cut into 250 μm-thick acute coronal slices. Up to five slices were then transferred to pre-incubation chamber filled with carbogenated recording ACSF composed of (in mM): 125 NaCl, 25 NaHCO_3_, 3 KCl, 1.2 Na_2_HPO_4_, 2 CaCl_2_, 2 MgCl_2_, 5 glucose, and 0.01 mg/l of Phenol Red (initial temperature: 32°C, cooled to room temperature) for at least one hour prior the recording.

#### 2.4.2. Multi-electrode array (MEA) recording

Experiments with the use of the MEA platform (Belle *et al*., 2021) were carried out at four daily time points on brainstem slices obtained from 23 rats (CD: n=11, HFD: n=12). Four to five slices were recorded at each time point in each diet. Slices were positioned in the recording wells of the MEA2100-System (Multichannel Systems GmbH, Germany) with the whole DMV localised upon the 6 × 10 recording array of a perforated MEA (60pMEA100/30iR-Ti, Multichannel Systems). Throughout the experiment, slices were perfused with fresh recording ACSF (2 ml/min), constantly bubbled with carbogen and heated to 32°C. Slices were allowed a minimum of 30 min to equilibrate before recordings were initiated. Data were acquired with MultiChannel Experimenter software (sampling frequency = 20 kHz; Multichannel Systems). Baseline recording was made for one hour. Then, drugs (orexin A, OXA 200 nM and GLP-1 1 µM) were diluted in 6 ml of fresh ACSF and bath applied in one hour intervals.

#### 2.4.3. Patch-clamp recording of DMV neurons

Tissue for patch-clamp experiments was isolated at two daily time points: at ZT3 for subsequent ‘late day’ (recording window: ZT6-12; CD: n=28, HFD: n=21 rats) and at ZT15 for subsequent ‘late night’ experiments (recording at ZT18-0, CD: n=27, HFD: n=14 rats). Following an incubation period, slices were placed in the recording chamber and constantly perfused (2 ml/min) with carbogenated recording ACFS heated to 32°C. For each recording, a single neuron in the borders of the DMV was randomly selected with a Zeiss Axioscope microscope fitted with infrared differential interference contrast under a 40x magnifying objective. Whole-cell configuration in both voltage- and current-clamp modes (VC and CC, respectively) was obtained with borosilicate glass pipettes produced with horizontal puller (resistance ∼7 MΩ; Sutter Instruments, USA) and suction applied by an Ez-gSEAL100B Pressure Controller (Neo Biosystem, USA). The signal was amplified by a SC 05LX (NPI, Germany) amplifier low-pass filtered at 2 kHz and digitised at 20 kHz. All stages of experiments were recorded with Signal and Spike2 (Cambridge Electronic Design Inc., UK) software. A liquid junction potential of −15 mV was added to the measured values of membrane potential.

Two intrapipette solutions with variant chloride concentration were used: normal intrapipette (nintra) containing (in mM): 125 potassium gluconate, 20 KCl, 10 HEPES, 2 MgCl_2_, 4 Na_2_ATP, 0.4 Na_3_GTP, 1 EGTA and 0.05% biocytin; or low chloride intrapipette solution (low Cl^-^intra) containing (in mM): 145 potassium gluconate, 10 HEPES, 2 MgCl_2_, 4 Na_2_ATP, 0.4 Na_3_GTP, 1 EGTA and 0.05% biocytin (in both cases: pH 7.4 adjusted with 5 M KOH; osmolality ∼300 mOsmol/kg). Low chloride concentration was introduced so as to shift the reversal potential for chloride current from −43 mV in nintra to −70 mV, in order to distinguish between inhibitory postsynaptic currents (IPSCs) visible as positive deflections and excitatory postsynaptic currents (EPSCs) seen as negative deflections. VC recordings were performed with nintra at holding potential of -65mV, whereas these using the lowCl^-^ intra at -50 mV. Negative rectangular pulses (duration: 1 s, amplitude: 25 mV) were applied every 60 s throughout the recording. CC recordings were exclusively carried out with the nintra with a holding current adjusted to set the membrane potential on -65 mV.

#### 2.4.4. Drugs

Orexin A (OXA; 200 nM, Bachem, Switzerland), glucagon-like peptide 1 (GLP-1; 1µM, Bachem), 6-cyano-7-nitroquinoxaline-2,3-dione (CNQX; 10 µM, Tocris, UK), DL-2-Amino-5-phosphonopentanoic acid (DL-AP5, 40 µM, Tocris), bicuculline methiodide (Bic, 20 μM, Tocris), tetrodotoxin citrate (TTX; 0.5 µM, Tocris) were stocked at 100x concentration at -20°C and were freshly diluted in the recording ACSF prior the application by bath perfusion.

#### 2.4.5. Post-recording immunostaining

To visualise the location of cells recorded near the anatomical borders of the DMV, slices used in patch clamp recordings were subsequently processed immunohistochemically and the recording site was determined. Briefly, following a successful recording, slices were fixed in 4% PFA diluted in PBS overnight in 4°C. Next, slices were rinsed twice in fresh PBS and placed in a permeabilising solution containing 0.6% TritX-100 (Sigma) and 10% NDS (Abcam) diluted in PBS for three hours in room temperature. Subsequently, sections were transferred to a solution containing Cy3-conjugated ExtrAvidin (1:250, Sigma) and primary antibodies against neuropeptide Y (NPY, raised in rabbit, 1:8000, Sigma) and incubated in 4°C overnight. Subsequently, slices were rinsed twice in PBS and incubated with anti-rabbit AlexaFluor 647-conjugated antisera (Jackson ImmunoResearch) for six hours at room temperature. Finally, slices were rinsed twice in PBS and mounted on glass slides in Fluoroshield™ (Sigma) and inspected under 10x magnification on an epifluorescence microscope (Axio Imager.M2, Zeiss). Biocytin-filled neurons were classified as DMV neurons based on their location in the NPY-ir-rich area as well as their large (>20 µm) cell body size (contrasting with the smaller neurons in the adjacent NTS).

#### 2.4.6. Spike-sorting and analysis of multi-electrode array data

Raw data were first exported to HDF5 files in Multi Channel DataManager (Multichannel Systems GmbH). Then, HDF5 files were remapped and converted to DAT format using a custom written MatLab script (R2018a version, MathWorks). Subsequently, DAT files were automatically spike-sorted with the use of KiloSort programme (Pachitariu *et al*., 2016) run in MatLab environment. To improve the speed of spike-sorting, a GPU was used (NVIDIA GeForce GTX 1050Ti GPU; CUDA 9.0 for Windows). Parallelly, raw data were also exported to CED-64 files with Multi Channel DataManager and further subjected to remapping and filtering with Butterworth band pass filter (fourth order) from 0.3 to 7.5 kHz by a custom-written Spike2 script. Subsequently, spike-sorted putative single units were transferred to CED-64 files using custom-made MatLab script and further refined in Spike2 (Spike2 8.11; Cambridge Electronic Design Ltd.) with the use of principal component analysis and autocorrelation.

Spontaneous neuronal activity was assessed in a 1800 s window, starting 30 min after the initiation of recording. Neuronal responses (activations and inhibitions evoked by the drug application) were calculated in 30 s bins with NeuroExplorer 6 (Nex Technologies, USA). A unit was classified as responsive if its single-unit activity varied by three standard deviations (SDs) from its baseline mean value. Amplitudes of these responses were calculated as a difference between maximal firing frequency (in 30 s bin) during the response and 10 minutes long mean baseline value.

#### 2.4.7. Analysis of patch-clamp data

Responsiveness to drug administration was measured with a custom-made script in MatLab (R2018a version, MathWorks). Changes in the whole-cell current were considered significant if they differed by more than three SDs from the averaged baseline values. Synaptic input was measured by manual selection of postsynaptic currents from 100 seconds of baseline recording with Mini Analysis Program (Synaptosoft, USA). Frequency was calculated for every condition. Additionally, current kinetics (rise time and decay time constant) were examined for neurons recorded with low Cl^-^intra (differentiated EPSC and IPSC). Curve fitting for decay time constant was performed for cells with minimal frequency of 0.08 Hz. Electrophysiological tests in current clamp mode were analysed with custom made Signal 5.07 scripts (Cambridge Electronic Design Ltd.).

### 2.5. Statistics

All statistical testing was performed in Prism 7 (GraphPad). Outliers were removed with the aid of ROUT (Robust regression and Outlier removal) method with coefficient Q = 0.05. Data were presented as individual values and mean. *P*<0.05 was deemed significant.

Two-way ANOVA was used to assess daily variation and dependence on dietary conditions in (1) the spontaneous DMV activity recorded on MEA, (2) frequency and kinetics of synaptic currents, (3) electrophysiological parameters of excitability and basic membrane properties and (4) responses to drug applications. Post-hoc multiple comparison was performed with Sidak’s test.

Fisher’s test was carried out to elucidate possible differences in the proportion of drug responsive neurons in the DMV between day and night.

## 3. RESULTS

### 3.1. Daily variation in neuronal activity of DMV neurons

The intracellular molecular clock drives SCN cells as well as neurons in extra-SCN oscillators including the NTS, to vary their spontaneous neuronal activity from day to night (Takahashi, 2017; Chrobok *et al*., 2020; Paul *et al*., 2020). To assess if DMV neurons also alter their firing rate across the day-night cycle, we used the MEA platform and recorded spontaneous multi-unit activity in 36 brainstem slices (17 from rats fed CD and 19 from animals fed the HFD; Fig. 1A,B) obtained from animals culled at one of four time points (ZT3, 9, 15 and 23; where ZT12 = lights-off). Recording was always initiated 2h following slice preparation. Using spike-sorting, we discriminated and identified putative single units in these DMV multi-unit activity records (Fig. 1C). Notable time of day variation in neuronal activity was detected (ZT: *p*<0.0001; two-way ANOVA), with the highest activity recorded at the end of light phase (ZT11). Thus, rat DMV neurons vary their spontaneous firing rate across 24 h, peaking at a similar late day phase as previously reported for the mouse NTS (Chrobok *et al*., 2020).

**Figure 1.**
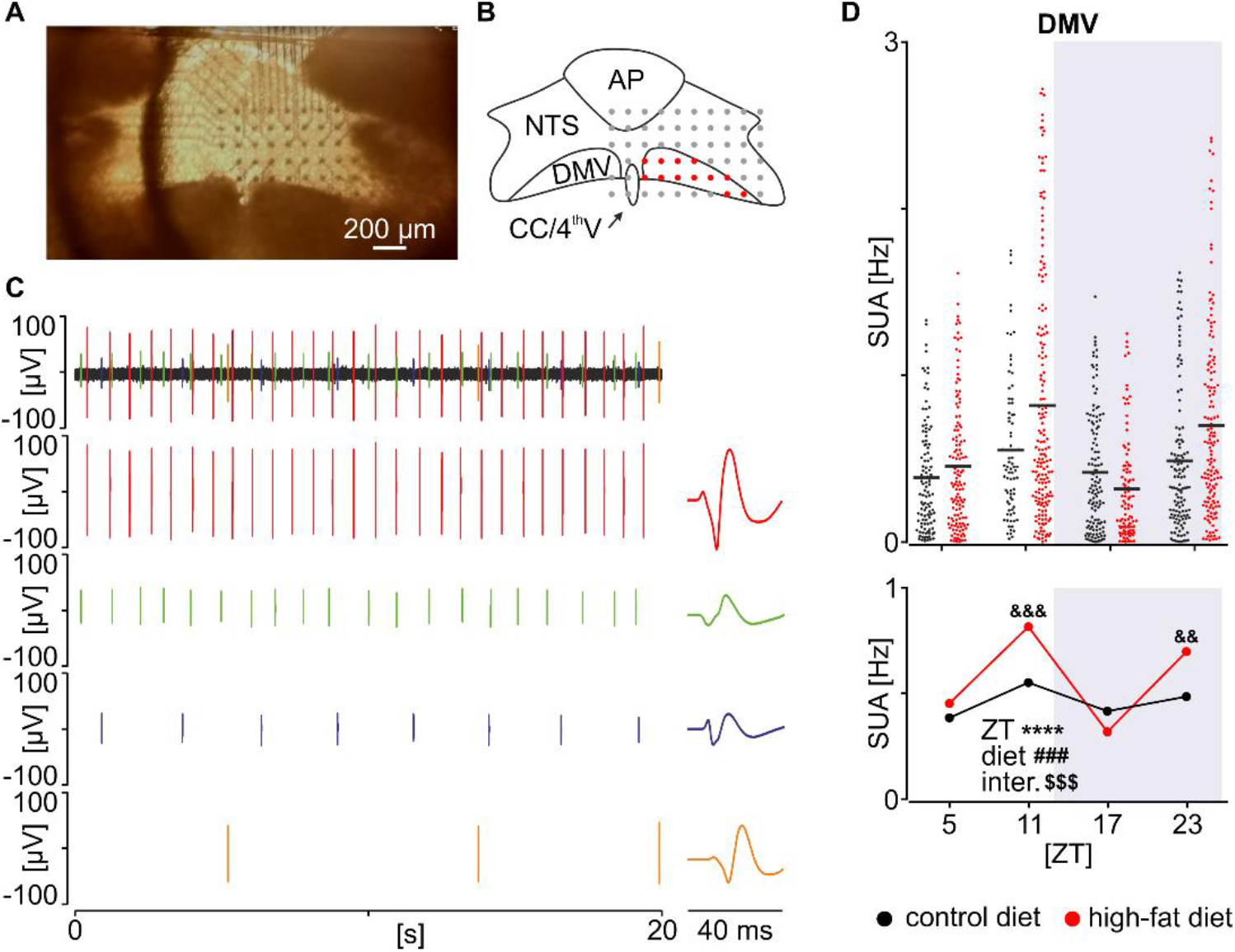
Daily variation in neuronal activity of dorsal motor nucleus of the vagus (DMV) is enhanced by short-term high fat diet (HFD). (**A**) Representative photomicrograph of the brainstem slice mounted on the multi-electrode array (MEA). (**B**) Outline of the dorsal vagal complex with recording locations reconstructed. Those localised to the DMV are coloured in red. AP – area postrema, CC/4^th^V – central canal/4^th^ ventricle, NTS – nucleus of the solitary tract. (**C**) Example 20 s recording trace with raw multi-unit extracellular signal (*top trace*) and four individual spike-sorted single units (*below*). Distinct average spike waveforms are shown for each single unit. (**D**) Scatterplot displaying all individual single-unit activity in the DMV (*above*) and a summary plot for the same data with means only (*below*). Data were plotted for four daily time points and two dietary conditions (****ZT *p*<0.0001, ###diet *p*=0.0005, $$$interaction *p*=0.0003, two way-ANOVAs; &&diet *p*=0.0015, &&&diet *p*=0.0003, Sidak’s multiple comparison test).

In mice, high-fat diet can eliminate circadian rhythmicity in NTS clock gene expression (Kaneko *et al*., 2009). Therefore, we next examined if diet affects the daily pattern in DMV neural firing. Short-term (two to three weeks) exposure to high-fat diet significantly altered this daily variation (*p*=0.0003; two-way ANOVA interaction) by elevating firing rate in the DMV (diet: *p*=0.0005; two-way ANOVA; Fig. 1D). Interestingly, post-hoc evaluation revealed this diet-evoked increase in neuronal activity to be significant only around the day-to-night (ZT11: *p*=0.0003) and night-to-day transitions (ZT23: *p*=0.0015, Sidak’s test; Fig. 1D). These findings suggest that daily changes in DMV neuronal firing are sensitive to dietary conditions. Statistical recapitulation of these results is presented in Supplementary Table 1.

### 3.2. Time of day variation in the synaptic input to the DMV

Daily variation in spontaneous neuronal activity may stem from daily change in intrinsic membrane properties, synaptic input, or a combination of these. To determine if DMV neurons are subject to day-night variation in synaptic input, we made patch-clamp recordings from these neurons at ZT6-12 (late day) or ZT18-0 (late night). First, we recorded all spontaneous postsynaptic currents (PSC) at -65 mV holding potential such that both excitatory (EPSC) and inhibitory currents (IPSC) were depicted as inward events (Fig. 2A,B). We found that the overall PSC frequency was higher at late day than late night (ZT: *p*=0.0443, two-way ANOVA; Fig. 2C,D). To precisely evaluate the polarity of these inputs [EPSCs (inward) vs IPSCs (outward)], we altered the [Cl^-^] of the intrapipette solution and made recordings at a holding potential of -50 mV (Fig. 2E). From these, we first identified that EPSCs constitute the vast majority of all PSCs in the DMV. Second, we observed a prominent temporal difference in EPSC frequency, with higher excitatory input at late day (ZT: *p*=0.0006, two-way ANOVA; Fig. 2F,G). Last, to determine if alterations in glutamate receptor responsiveness contributed to the postsynaptic mechanisms shaping responses to synaptic input, we measured EPSC kinetics such as their rise and decay time. No day-to-night differences in EPSC rise time (ZT: *p*=0.0783; Fig. 2H) or decay time constant (ZT: *p*=0.3050, two-way ANOVAs; Fig. 2I) were detected. This indicates that daily changes in presynaptic mechanisms are predominately responsible for enhanced excitatory input to DMV neurons at the late day phase.

**Figure 2.**
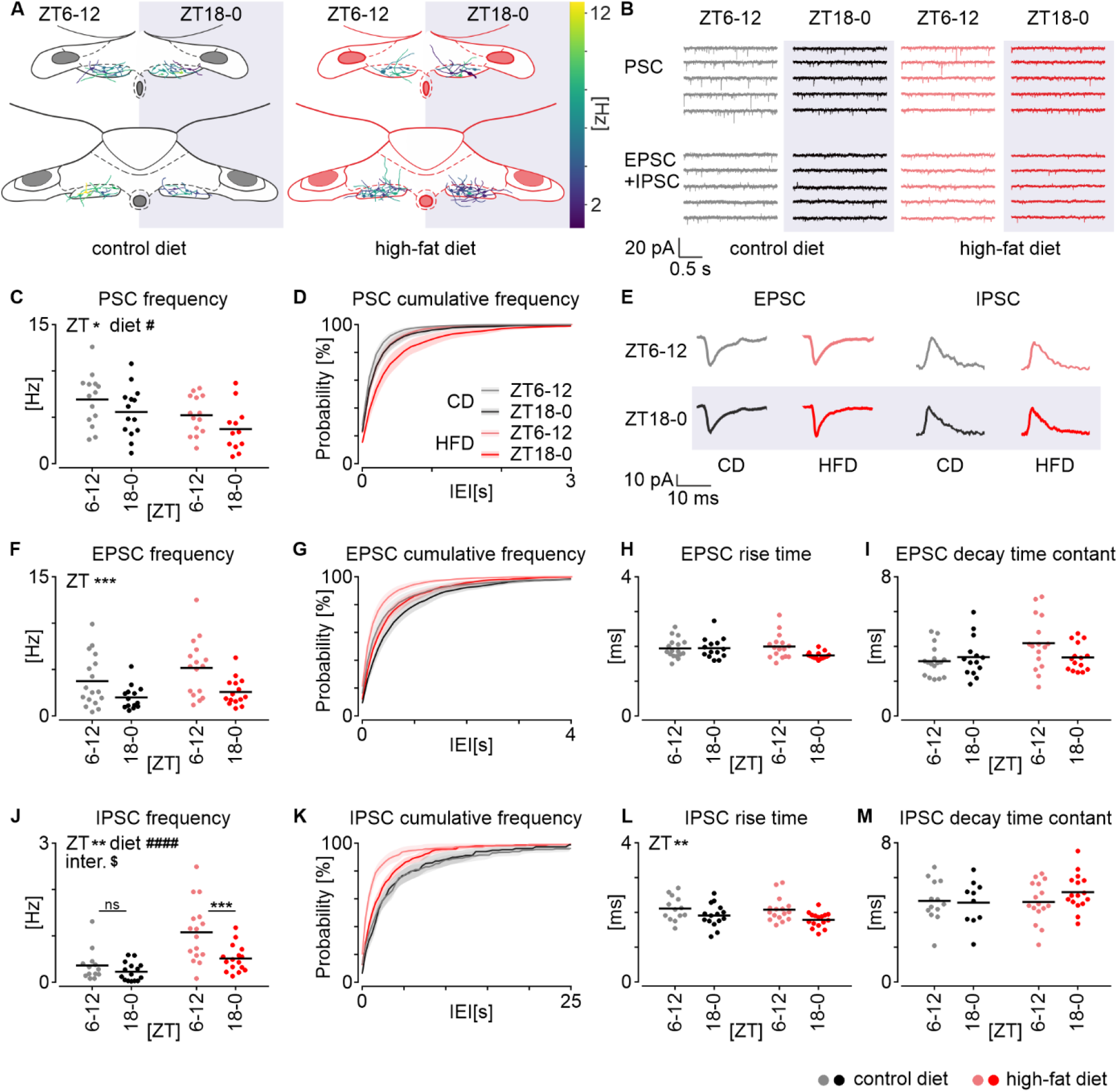
Spontaneous synaptic input to dorsal motor nucleus of the vagus (DMV) displays robust day-to-night variation which is altered by high fat diet (HFD). (**A**) Schematic location of DMV neurons superimposed on two planes, recorded at late day (ZT6-12) or late night (ZT18-0) from slices obtained from HFD and control diet (CD) rats. Warm colours code higher total frequency of postsynaptic currents (PSC). (**B**) Representative examples of synaptic activity throughout 10 s, for each group of rats. EPSC – excitatory postsynaptic currents, IPSC – inhibitory postsynaptic currents. (**C**) Daily changes in the frequency of total synaptic input to the DMV (ZT: \**p*=0.0443, diet: #*p*=0.0137) with corresponding cumulative frequency plot in **D** (bin=100 ms). (**E**) Example EPSC and IPSC for all experimental groups depicted as an average of all events from one neuron. (**F**) Day-to-night difference in the frequency of EPSC (ZT: \*\*\**p*=0.0006) with cumulative frequency traces shown in **G** (bin=100 ms). (**H**). Length of an averaged EPSC rise time. (**I**) Fitted EPSC decay time constant. (**J**) Daily changes in the IPSC frequency (ZT: \*\**p*=0.0018, diet: ####*p*<0.0001, interaction: $*p*=0.0472). Note that significant changes were noted in HFD rats (\*\*\**p*=0.0005), but not CD (ns *p*=0.6321). Corresponding cumulative frequency plot is shown in **K** (bin=400 ms). (**L**) Daily change in the average IPSC rise time (\*\**p*=0.0035). (**M**) Fitted IPSC decay time constant. All statistical testing was performed with ordinary two-way ANOVA followed by Sidak’s multiple comparison test (results drawn above black bars). In all plots, grey codes recordings at late day from rats fed CD, black – at late night from CD, pink – at late day from HFD and red – at late night from HFD.

Subsequently, we evaluated inhibitory synaptic inputs to the DMV and found a significant time of day effect on IPSC frequency (ZT: *p*=0.0018; Fig. 2J,K). Interestingly, this daily change in inhibitory inputs was also accompanied by a change in postsynaptic mechanisms as the IPSC rise time was significantly elongated at ZT6-12 (ZT: *p*=0.0035; Fig. 2L), while the decay did not vary from late day to night (ZT: *p*=0.4746, two-way ANOVAs; Fig. 2M). These findings indicate that both pre- and postsynaptic mechanisms contribute to elevated inhibitory tone on DMV neurons at late day.

Finally, to evaluate if diet influenced this day-night change in synaptic inputs to DMV neurons, we compared recordings from CD and HFD rats. We found that high-fat diet reduced the frequency of all PSCs (recorded at -65 mV) (diet: *p*=0.0137, two-way ANOVA; Fig. 2C,D), but did not affect the frequency of EPSCs (recorded with the altered intrapipette solution at -50 mv; diet: *p*=0.0974, two-way ANOVA; Fig. 2F,G). However, assessment of inhibitory synaptic input to DMV neurons indicated that high-fat diet increased IPSC frequency (diet: *p*<0.0001, two-way ANOVA; Fig. 2J,K). Additionally, diet had a significant effect on the daily variation in IPSC frequency (diet: *p*=0.0472, two-way ANOVA interaction; Fig. 2J) with the frequency of inhibitory synaptic input significantly elevated at late day in HFD (*p*=0.0005), but not CD fed rats (*p*=0.6321, Sidak’s test; Fig. 2J). Thus, these results suggest that high-fat diet selectively increases late day inhibitory input to DMV neurons, without affecting the excitatory input to these cells. Moreover, no diet-related alterations of EPSC (rise time, diet: *p*=0.2759, decay time constant, diet: *p*=0.0803; Fig. 2H,I) or IPSC kinetics were observed (rise time, diet: *p*=0.3498, decay time constant, diet: *p*=0.3897, two-way ANOVAs; Fig. 2L,M). Thus, these effects of diet on the DMV synaptic activity are attributable to changes at presynaptic sites. All statistics summarising the synaptic activity section are presented in Supplementary Table 2.

### 3.3. Day to night change in intrinsic excitability of DMV neurons

For increased detection and responsiveness to afferent information, neurons increase their resting membrane potential such that a small change in excitatory input can trigger action potential generation. Alternatively, without such a change in resting membrane potential, they can alter voltage-dependent mechanisms (e.g. increase membrane resistance) to elevate their excitability (Hille, 1992). To evaluate if DMV neurons adjust their excitability to changing synaptic input in a time of day fashion, we made patch clamp recordings *ex vivo* at late day and late night on hindbrain slices obtained from CD and HFD rats. First, high amplitude (1 nA) current ramp stimulation lasting 1 s was applied on DMV neurons manually held just below the threshold of action potential generation, at membrane potential of -65 mV (Fig. 3A). Robust time of day related change in the response to this stimulation was observed: at late day DMV neurons exhibited higher maximal instantaneous discharge frequencies as a result of ramp-evoked depolarisation (ZT: *p*<0.0001; Fig. 3B) and overall generated a substantially higher number of action potentials during the stimulation (ZT: *p*<0.0001; Fig. 3C). DMV neurons of HFD rats showed heightened responsiveness to ramp stimulation with higher maximal instantaneous firing frequency (diet: *p*=0.0252; Fig. 3B) and increased total number of action potentials evoked by current injection (diet: *p*=0.0103, two-way ANOVAs; Fig. 3C), compared to CD.

**Figure 3.**
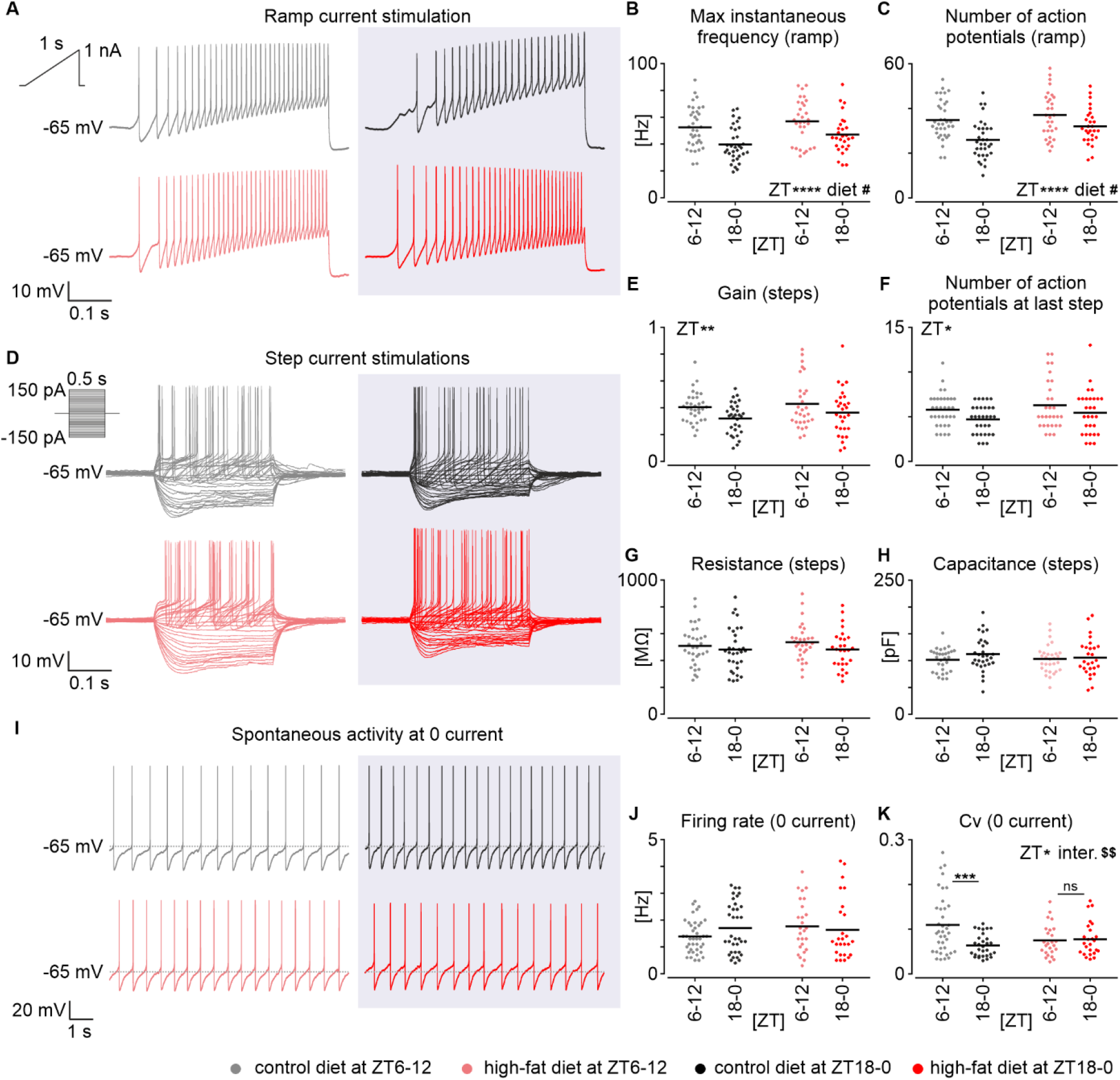
Late day increase in the excitability of the dorsal motor nucleus of the vagus (DMV). (**A**) Example traces of ramp current stimulation recorded from single DMV neurons at late day and late night in slices obtained from rats fed high fat (HFD) or control diet (CD). (**B**) Apparent decrease in the maximal instantaneous frequency elicited on the current ramp at late night (\*\*\*\**p*<0.0001) and its significant reduction under high-fat diet (#*p*=0.0252). (**C**) Corresponding effect in the number of action potential generated during the ramp (ZT: \*\*\*\**p*<0.0001, diet: #*p*=0.0103). (**D**) Representative traces of step current protocol applied on the DMV neurons from the same four experimental groups. (**E**) Evident rise in the excitability curve slope (gain) at late day comparing to late night (\*\**p*=0.0065). (**F**) Parallel diurnal rise in the number of action potentials fired at the last step of the stimulation protocol (\**p*=0.0191). (**G&H**) Basic membrane properties of DMV neurons: membrane resistance and capacitance. (**I**) Representative spontaneous firing activity of DMV neurons at holding current 0. (**J**) Spontaneous firing rate measured over 10 s. (**K**) Regularity of single DMV neurons calculated as the coefficient of variation (Cv) of inter-spike intervals. Daily variation (\**p*=0.0140) explained by a loss of regularity at late day in the CD-fed rats only (interaction: $$*p*=0.0061, CD: \*\*\**p*=0.0002, HFD: ns *p*=0.9754). All statistical testing was performed with ordinary two-way ANOVA followed by Sidak’s multiple comparison test (results drawn above black bars). In all plots, grey codes recordings at late day from rats fed CD, black – at late night from CD, pink – at late day from HFD and red – at late night from HFD.

Next, we used a step current stimulation protocol comprising 30 steps from -150 to +150 pA (every 10 pA, duration: 0.5 s; Fig. 3D) to generate excitability curves and characterise possible changes in fundamental electrophysiological properties (such as membrane resistance and capacitance) of DMV neurons. Again, DMV neurons exhibited significantly higher excitability at ZT6-12 comparing to ZT18-0, measured as higher excitability curve slopes (gain, ZT: *p*=0.0065; Fig. 3E) resulting from increasingly greater numbers of evoked action potentials at consecutive steps (at last step, ZT: *p*=0.0191; Fig. 3F). In this step protocol, we did not observe any diet-related effects (gain, diet: *p*=0.2033, action potentials at last step, diet: *p*=0.1288, two-way ANOVAs; Fig. 3E,F). Since this step protocol did not depolarise DMV neurons to the same extent as the ramp stimulation did, this suggests that diet-evoked effects are voltage dependent. This assertion is further supported by the results of measurements of passive membrane properties with the hyperpolarising step current injections, which did not vary from late day to late night (resistance, ZT: *p*=0.1314, capacitance, ZT: *p*=0.2111; Fig. 3G,H) or between diets (resistance, diet: *p*=0.5995, capacitance, diet: *p*=0.6255, two-way ANOVAs; Fig. 3G,H).

Finally, we recorded spontaneous activity of DMV neurons in these four groups of animals in current clamp mode at holding current = 0 (Fig. 3I). The majority of recorded cells (∼85 %) exhibited spontaneous, regular, low frequency firing around 1-2 Hz (as previously reported; Browning *et al*., 1999), with no evident differences in discharge rate among groups (ZT: *p*=0.5674, diet: *p*=0.3379, two-way ANOVA; Fig. 3J). Additionally, the coefficient of variation in action potential generation (a measure of spiking regularity) was assessed and found to be subject to time of day variation (Cv, ZT: *p*=0.0140, two-way ANOVA; Fig. 3K), with neurons at late day firing less regularly. This variation in firing regularity was detected under CD, but lost under HFD conditions (*p*=0.0061, two-way ANOVA interaction; CD: *p*=0.0002, HFD: *p*=0.9754; Sidak’s tests; Fig. 3K), with HFD DMV neurons exhibiting regular firing at both late day and late night phases. Statistical details regarding this dataset can be found in Supplementary Table 3. These results provide evidence for increased excitability of DMV neurons at late day, coincident with the time of a day at which there is elevated synaptic input to these neurons. Additionally, they show that high-fat diet increased their excitability and regularity of firing at this late day phase. Further, these investigations suggest voltage-dependency of ionic mechanisms underlying daily and diet-evoked changes in neuronal excitability in the DMV.

### 3.4. Responsiveness to neuromodulators in the DMV across 24 h

Activity of DMV neurons is not only shaped by fast glutamate and GABA synaptic transmission, but is also influenced by neuromodulators conveying information on metabolic and arousal state (Travagli & Anselmi, 2016). To examine this, we investigated the responsiveness of DMV neurons to two peptides implicated in opposing physioogical states: the preprandial orexin A (OXA) that stimulates gastric function (Krowicki *et al*., 2002), and postprandial GLP-1 which promotes gastroinhibition (Holmes *et al*., 2009). Initially, we used the MEA platform and tested DMV neuronal responsiveness to these neuropeptides at four time points over 24 h in CD and HFD rats. The majority of single units recorded in the DMV were responsive to both OXA (200 nM; Fig. 4A) and GLP-1 (1 µM; Fig. 4D), predominately increasing their firing rate to both. However, using this experimental approach, we did not note any significant effects of time of day or diet on the amplitude of the response to OXA (ZT: *p*=0.3337, diet: *p*=0.8543, interaction: *p*=0.3013; Fig. 4B) or GLP-1 (ZT: *p*=0.8256, diet: *p*=0.1012, interaction: *p*=0.1498, two-way ANOVAs; Fig. 4E). For more statistical details on this dataset see Supplementary Table 4. These extracellular recordings suggest that DMV neurons are responsive to metabolically relevant neuropeptides across the 24 h cycle, but with no obvious day-to-night change in magnitude of response.

**Figure 4.**
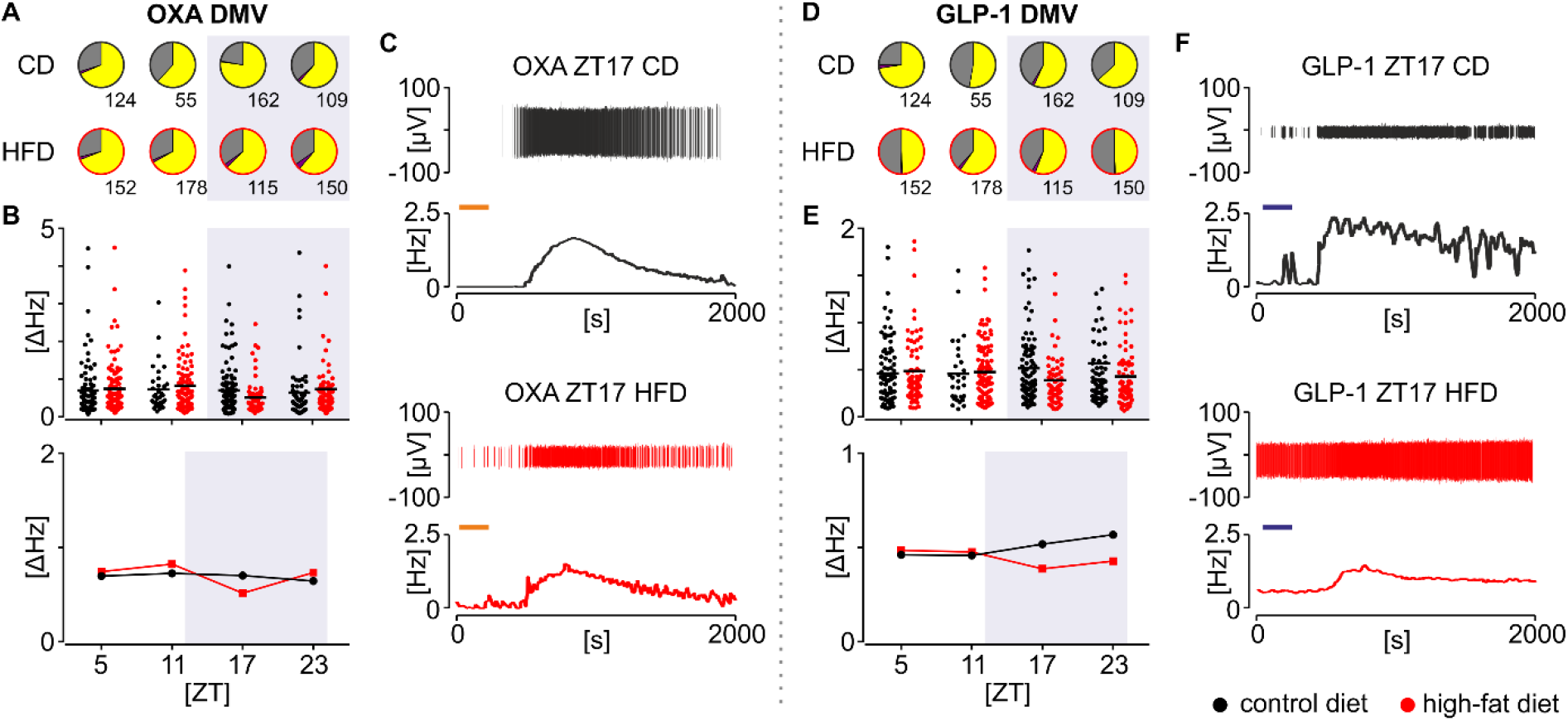
Responsiveness to orexin A (OXA; 200 nM) and glucagon-like peptide-1 (GLP-1; 1 μM) in the dorsal motor nucleus of the vagus (DMV) across 24 h. (**A & D**) Proportion of single units excited (yellow), suppressed (purple) and non-sensitive (grey) to drug application at four daily time points. Black outline of the pie chart codes control diet (CD), whereas red depicts high fat diet (HFD). Numbers code total ns of neurons tested. (**B & E**) Scatterplots displaying individual amplitudes of excitatory drug responses (*above*) and a summary plot for the same data with means only (*below*). (**C & F**) Representative excitatory responses to drug application, presented as spike-sorted single unit wave marks (*above*) and firing frequency histograms (*below*). Bin = 30 s. In all subpanels black codes CD and red – HFD.

### 3.5. Electrophysiological responses to metabolic signals exhibit day-to-night variation in the DMV, disturbed by high fat diet

In the extracellular recording configuration, neuronal membrane potential cannot be controlled and therefore the possible contribution of voltage-dependent factors to the response of neurons to neuromodulators cannot be distinguished. Therefore, we used voltage clamp recordings to investigate with greater precision the effects of diet (CD vs HFD) and time-of-day influences (ZT6-12 or ZT18-0) on the electrophysiological responses of DMV neurons to orexin A (OXA, 200 nM) and GLP-1 (1 µM). Since diet-evoked changes in DMV electrophysiological parameters exhibited voltage-dependency (Fig. 3), we next tested the response of DMV neurons to OXA and GLP-1 in voltage clamp configuration at two different holding potentials (−50 and -65 mV). The -65 mV holding potential was used to mimic the endogenous hyperpolarised state of DMV neurons, whereas -50 mV mimics their depolarised state, near their threshold of action potential generation.

First, OXA was administered at holding potential of -50 mV in standard artificial cerebro-spinal fluid (ACSF) (Fig. 5A). The amplitude of OXA-evoked current measured at late day was significantly larger than that elicited at late night (ZT: *p*=0.0015), but no overall effect of diet on responses to OXA was detected (diet: *p*=0.9842, two-way ANOVA; Fig. 5C). In more hyperpolarised conditions (−65 mV), there were no time of day differences in the current evoked by OXA (ZT: *p*=0.6383), but high-fat diet significantly blunted responses to this neuropeptide (diet: *p*=0.0176, two-way ANOVA; Fig. 5B,D). A total of 132 DMV neurons were tested at one of the two holding potentials in the standard ACSF with similarly high proportion of cells responding to OXA in HFD (late day: 91%, late night: 96%, *p*>0.9999) and CD rats (late day: 83%, late night: 82%, *p*=0.6233, Fisher’s tests; Fig. 5E). To evaluate whether the response to OXA was generated intrinsically, DMV neurons were tested at -65 mV for response to this neuropeptide in the presence of tetrodotoxin (TTX, 0.5 µM) and fast synaptic transmission blockers (DL-AP5 40 µM, CNQX 10 µM, bicuculline 20 µM) in the ACSF. Under these synaptic isolation conditions, OXA application evoked inward current from all DMV neurons examined (41/41; Fig. 5B), indicating that this neuromodulator acts postsynaptically to alter DMV neuronal activity. Thus DMV neurons alter their responsiveness to OXA from late day to late night, with this effect being dependent on membrane potential and blunted by high-fat diet.

**Figure 5.**
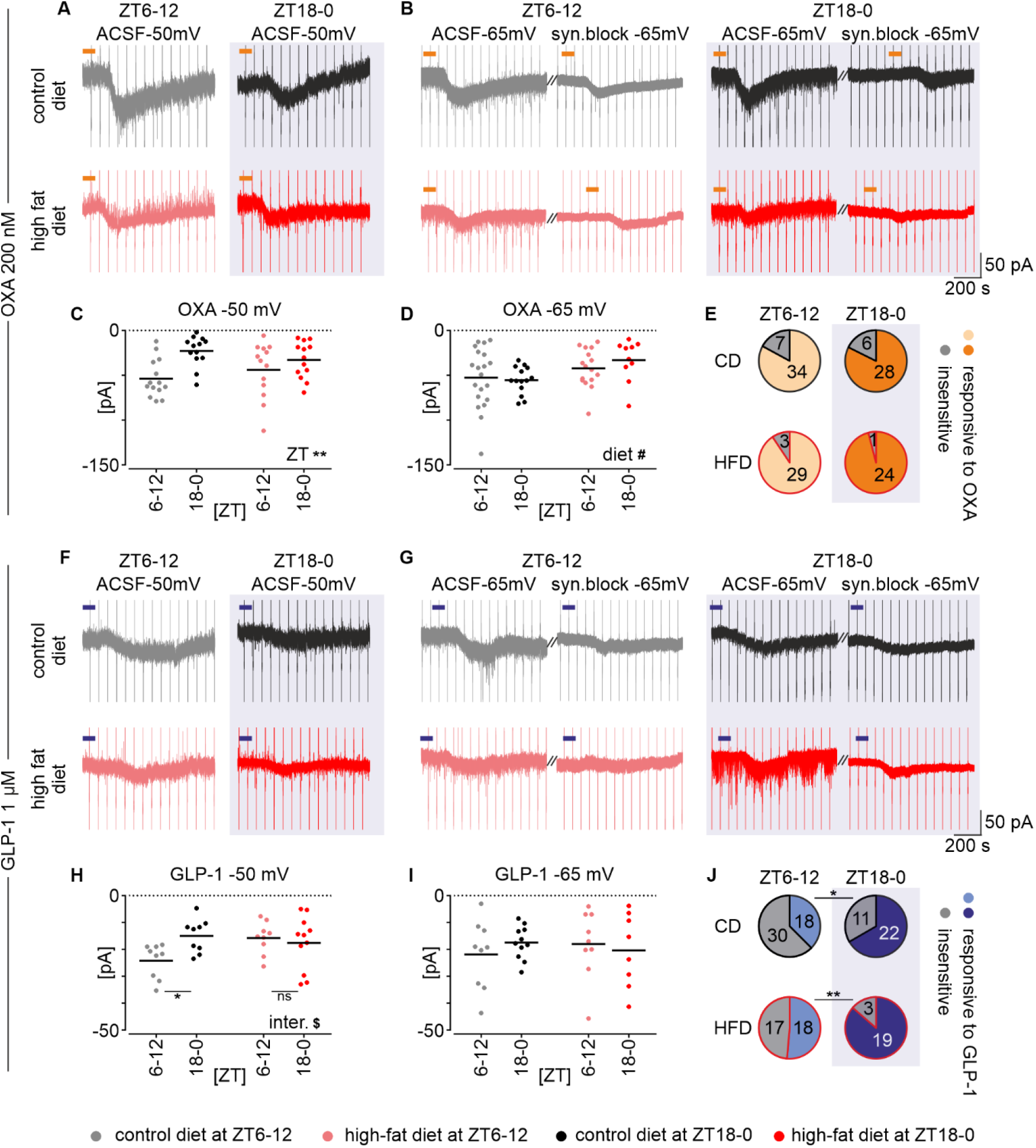
Day-night variation in the responsiveness of neurons in the dorsal motor nucleus of the vagus (DMV) to orexin A (OXA) and glucagon-like peptide 1 (GLP-1). (**A**) Example raw voltage clamp recording traces (holding potential = -50 mV) of DMV neurons from late day (ZT6-12) or late night (ZT18-0) obtained from rats fed high fat (HFD) or control diet (CD). (**B**) Representative recordings at - 65 mV in standard artificial cerebrospinal fluid (ACSF) or ACSF containing tetrodotoxin (0.5 μM) and fast synaptic blockers: DL-AP5 (40 µM), CNQX (10 µM) and bicuculline (20 µM) (syn. block). Orange bars pinpoint bath applications of OXA (200 nM). (**C**) Current amplitude in response to OXA at -50 mV with a clear daily variation (\*\**p*=0.0015). (**D**) Inward current evoked by OXA at -65 mV (diet: #*p*=0.0176). (**E**) Number of responsive DMV neurons tested in each of four experimental groups. (**F**) Example traces at -50 mV showing responses to GLP-1 (blue bars) of DMV neurons. (**G**) Representative recordings presenting current responses to GLP-1 at -65 mV, in control ACSF or ACSF containing tetrodotoxin and blockers of fast synaptic transmission (syn. block). (**H**) Late day rise in the amplitude of current evoked by GLP-1 application at -50 mV in rats fed CD only (interaction: $*p*=0.0306, CD: \**p*=0.0272, HFD: ns *p*=0.8323). (**I**) No changes in current amplitudes evoked by GLP-1 application at -65 mV. (**J**) Nocturnal rise in the proportion of GLP-1-responsive neurons in the DMV in both CD (\**p*=0.0131) and HFD rats (\*\**p*=0.0100). All current amplitudes were compared using two-way ANOVA followed by Sidak’s multiple comparison test (results drawn above black bars). Response occurrence ratios were tested with Fisher’s test. In all plots, grey codes recordings at late day from rats fed CD, black – at late night from CD, pink – at late day from HFD and red – at late night from HFD.

Next, we tested time of day and diet-dependent actions of GLP-1 on the whole cell current of DMV neurons (Fig. 5F,G). Similarly to OXA, GLP-1 applied at -50 mV evoked inward currents of a higher amplitude at ZT6-12 comparing to ZT18-0, but only in rats fed CD (*p*=0.0306, two-way ANOVA interaction; CD: *p*=0.0272, Sidak’s test; Fig. 5H). The late day increase in GLP-1-evoked current amplitude was abolished by high-fat diet (HFD: *p*=0.8323; Sidak’s test; Fig. 5H). Similar to OXA, these time of day differences were not detected at lower holding potential (−65 mV; *p*=0.7957). In contrast to OXA, no diet-dependent changes were noted in the GLP-1 action at -65 mV (diet: *p*=0.8955, two-way ANOVA; Fig. 5I). In total, responsiveness of 138 DMV neurons to GLP-1 was tested in standard ACSF at both potentials, with a significantly higher percentage of neurons responsive at late night for rats fed CD (38% vs. 67%, *p*=0.0131) or HFD (51% vs. 86%, *p*=0.0100, Fisher’s tests; Fig. 5J). Subsequent assessment of these GLP-1 responsive DMV neurons in the presence of TTX (0.5 µM) and fast synaptic blockers (DL-AP5 40 µM, CNQX 10 µM, bicuculline 20 µM) revealed similar changes in current (19/19 neurons tested; Fig 5G). This implicates a direct postsynaptic action of GLP-1 on DMV neurons. Statistical summary for these sections is depicted in Supplementary Table 5.

These observations indicate that responsiveness of DMV neurons to metabolically relevant neuromodulators is elevated at the late day phase and that this is blunted by high-fat diet. These findings add further credence to the hypothesis that voltage-dependent mechanisms underpin daily rhythmicity of the DMV.

## 4. DISCUSSION

Here we report for the first time daily variation in synaptic events and neuronal activity in the DMV, a brain structure of key importance to the parasympathetic system. Our findings indicate that compared to the late night, DMV neurons increase their firing rates and excitability at late day, which is accompanied by a significant rise in the frequency of predominately excitatory synaptic input. These are paralleled by elevated late day responsiveness to OXA and GLP-1, neuromodulators implicated in the control of food intake. Unexpectedly, short-term consumption of a high-fat diet increased DMV neuronal firing and excitability, while blunting their responses to these periprandial cues. These findings indicate complex interactions of diet and time of day on DMV neuronal activity.

Previous investigations have determined that the DVC exhibits daily and circadian variation in molecular and neuronal activities (Herichová *et al*., 2007; Kaneko *et al*., 2009; Ubaldo-Reyes *et al*., 2017; Chrobok *et al*., 2020). Through long-term bioluminescence imaging, we found that the AP was the most robust circadian oscillator amongst structures of the DVC, and contained a high density of *Per2*-expressing clock cells. At the same time, a target of AP efferents, the NTS, (Shapiro & Miselis, 1985; Hay & Bishop, 1991) rhythmically expressed a lower density of *Per2* cells for up to a week *ex vivo*. In that analysis, the DMV was the least robust circadian oscillator among these three – *Per2* expression in slice culture was expressed only transiently and diminished after one 24 h cycle in culture (Chrobok *et al*., 2020). To our knowledge, circadian rhythmicity of efferent vagal activity has not yet been reported, but the afferent branch of vagus nerve exhibits daily and circadian properties. The somas of sensory vagal neurons are localised in nodose ganglia and these demonstrate rhythmic clock gene expression as well as a daytime (ZT6-9) peak in mechanosensitivity (Kentish *et al*., 2013, 2016, 2019). These observations raise the possibility that daily variation in DMV molecular and neuronal activity is dependent on recurrent circadian input from other DVC structures as well as signals from the peripheral nervous system.

To communicate its circadian phase information to the rest of the body, the AP/NTS must control the DMV and both *in vivo* as well as in *ex vivo* brain slice preparations, DMV neuronal firing is indeed under control of the AP and NTS (Morest, 1967; Shapiro & Miselis, 1985; Davis *et al*., 2004). It has been unclear as to whether the DMV, which directly controls vagal tone, exhibits time of day variation in its neuronal activity. In our previous MEA study on the mouse DVC, it was not possible to reliably delineate the DMV from the suprajacent NTS. Here, the larger anatomy of the rat DVC enabled us to distinguish MEA electrodes in the DMV from those in the adjacent areas. In contrast to overt day-to-night variation in neuronal activity of the rat NTS (reported by our parallel study (Chrobok *et al*., 2021*b*)), here we observe that the rat DMV demonstrates low amplitude time of day variation in neuronal firing is consistent with the low level of clock gene expression observed in this structure (Chrobok *et al*., 2020). Alternatively, this daily change in neuronal activity may be a fingerprint of the upstream NTS, whose firing rates exhibit profound day-to-night variation in the same brain slice preparation (Chrobok *et al*., 2021*b*). Overall, this 24 h variation in the DMV firing likely originates from multiple sources, including a combination of intrinsic drive of DMV neurons to alter firing rate and daily change in synaptic input to the DMV.

To attempt to differentiate these contributions, we performed patch clamp recordings to reveal more subtle characteristics in DMV electrophysiology. With these, we identified extrinsic and intrinsic sources for time of day variation in DMV function. First, spontaneous synaptic input was elevated at late day which was due to increased EPSCs. Although we did not identify the source of this synaptic input, it could reflect circadian changes in vagal afferents since these signals to the DMV are predominately excitatory (Raab & Neuhuber, 2007). The NTS innervates the DMV, but the polarity of this input is mostly inhibitory (Davis *et al*., 2004) and since the IPSC frequency of DMV neurons did not robustly change from day to night in control conditions, then circadian activity in the NTS is not likely to be communicated to the DMV in this way. However, since the kinetics of DMV IPSCs showed time of day variation, then this suggests that these neurons can intrinsically alter how they process inhibitory input. Such postsynaptic changes could originate from several factors including alteration in the phosphorylation state of GABA receptor subunits, channel density, subunit composition or their subcellular localisation (Krishek *et al*., 1994; Browne *et al*., 2001).

Second, we found a late day rise in DMV neuronal excitability that occurs in parallel with elevated synaptic input. This increase is manifested in higher maximal instantaneous firing frequency (lower inter-spike interval) and higher gain in DMV neuronal firing. Thus, intrinsic mechanisms of these neurons enable the generation of higher firing rates in response to depolarising stimuli at the late day phase. The magnitude of these day-night changes was more distinct at elevated membrane depolarisation, suggesting a voltage-dependent mechanism. We speculate that these changes arise from the late day downregulation of distinct potassium conductances. A candidate here is the transient A-type potassium current (*I*_*A*_), which increases the delay to next spike (Luther & Tasker, 2000; Baranauskas, 2007). Indeed our observation of the late day decrease in regularity of action potential firing by DMV neurons is consistent with this idea since regularity of neuronal firing is typically guarded by *I*_*A*_ (Khaliq & Bean, 2008). It is notable that in the dorsal SCN, expression of *I*_*A*_ channel subunits varies in a circadian fashion, peaking during the day (Itri *et al*., 2010). Further research is required to test whether *I*_*A*_ contributes similarly to daily variation in the regularity of DMV firing rate.

Neurons in the SCN, NTS, and ventrolateral geniculate nucleus exhibit time of day adjustments in their responsiveness to neuromodulators (Belle & Piggins, 2017; Chrobok *et al*., 2020, 2021*a*). Here, using patch clamp to control for membrane potential, we found DMV neurons to exhibit daily changes in responsiveness to OXA and GLP-1 (peptides implicated in the control of food intake). Moreover, these higher amplitude responses were noted at the time of increased excitability and synaptic input to these DMV neurons. Therefore, this temporally dependent accentuated sensitivity to peptides with opposing effects on gastric motility (Krowicki *et al*., 2002; Holmes *et al*., 2009), suggests a nonspecific preparedness for processing metabolic information at this time of day. Interestingly, these day-to-night differences in DMV neurons responsiveness to OXA and GLP-1 were diminished at more hyperpolarised membrane potentials, paralleling our observation on the voltage-dependence of daily changes in excitability of these DMV cells. Further, our MEA recordings revealed the DMV to be sensitive to these metabolic neuropeptides across the 24 h cycle. However, with the extracellular recording configuration, we did not detect any daily variation in the amplitude of these responses and this may be due to diverse baseline firing among neurons in different time points and voltage-dependence of ionic mechanisms underlying studied excitations (Acuna-Goycolea & van den Pol, 2004; Kukkonen, 2016).

The parameters of diet schedule as well as the age at which the diet is consumed influence vagal neurocircuitry. For instance, perinatal high-fat diet can increase inhibition of DMV neurons (McMenamin *et al*., 2018), while an acute 3-5 days of consuming high-fat diet in adult rats upregulates glutamatergic signalling to the DMV (Clyburn *et al*., 2018). Excess calorie consumption can also disrupt circadian timekeeping in the DVC. In adult mice, long-term consumption of high-fat diet eliminates rhythmic clock gene expression in the NTS (Kaneko *et al*., 2009). In our parallel study, we also found that short-term high-fat diet reduces neuronal activity of the NTS and blunts its daily rhythm (Chrobok *et al*., 2021*b*). As the DMV activity heavily depends on the NTS input, it is most likely that malfunctioning of this upstream DVC area under high-fat diet influences neuronal firing of the DMV. Diet can further alter the afferent branch of the vagal circuit. Consumption of a high-fat diet can abrogate circadian variation in vagal mechanosensitivity without altering clock gene expression in nodose ganglia (Kentish *et al*., 2016). Here we show, that high-fat diet unexpectedly boosted daily variation in neuronal activity of the DMV, increasing late day firing. We hypothesise, that this increase of neuronal firing in the DMV may be attributed to the disinhibition from a predominately inhibitory NTS, whose activity is heavily impaired by high-fat diet, particularly at late day (Chrobok *et al*., 2021*b*). Interestingly, the increased neuronal activity in the DMV by high-fat diet was not observed when recorded in the long-term set up (Chrobok *et al*., 2021*b*); in contrast to the acute ones, those longer-term (∼30 h) registrations allow to assess circadian changes in neuronal activity without any inputs caused by animal behaviour preceding the cull (such as feeding). Thus, the elevated DMV firing under high-fat diet is most likely a combination of disturbances intrinsic to the DVC and altered patterning of food intake.

Additionally after 2-3 weeks of diet, we observed a slight decrease in synaptic input to DMV, without any significant changes in EPSC frequency. Thus, due to the polarity of these changes, we speculate that they are unlikely to arise from afferent vagal activities. Since high-fat diet abolishes clock gene expression in the mouse NTS (Kaneko *et al*., 2009), it is tempting to speculate that such dietary conditions reduce tonic inhibitory drive from NTS to DMV. However, as high-fat diet elevated IPSCs from DMV neurons, then the increase in DMV neuronal activity is unlikely to be attributable to the influence of diet on inhibitory afferents to the DMV. Alternatively, high-fat diet may enhance the intrinsic excitability of DMV neurons and this is what we observed in this study. Interestingly, perinatal high-fat diet is reported to decrease DMV excitability (Bhagat *et al*., 2015), suggesting that effects of diet on vagal neurocircuitry varies markedly at a different stages of the lifespan.

Previous studies report that perinatal and adult high-fat diet impair DMV responsiveness to the neuromodulators, cholecystokinin and GLP-1 (Bhagat *et al*., 2015). However, in this investigation, we found that this calorie dense diet unequivocally suppresses responsiveness of vagal motoneurons to OXA and GLP-1. Further we observed that high-fat diet eliminates daily variation in the amplitude of DMV neuronal response to exogenous GLP-1. Diet-related changes in the modulation of the DMV by metabolically relevant peptides are not limited to brainstem, but can also act upstream at the lateral hypothalamic source of DMV afferents, as high-fat diet reduces orexin synthesis in the lateral hypothalamus (Lin *et al*., 2000; Nobunaga *et al*., 2014).

In summary, these findings indicate that synaptic input and intrinsic neuronal activity in the DMV can vary with time of day and are influenced by diet. The increase in activity and responsiveness to neuromodulators at the late day phase may function to prepare the DMV for processing the nocturnal rise in central and peripheral ingestive signals. There are potentially multiple sources underpinning these temporal changes including (1) intrinsic changes in DMV excitability and responsiveness to neuromodulators, (2) extrinsic variation in synaptic input, and (3) both intrinsically and extrinsically regulated alterations in DMV neuronal firing. Additionally, here we observed that short-term exposure to high-fat diet unexpectedly elevates the activity of DMV neurons, while blunting their responsiveness to neuromodulators. The function(s) of these daily changes in the DMV are not yet defined, but potentially contribute to temporal alteration in parasympathetic tone. Thus, our study is first to report possible daily differences in the efferent branch of the vagal system.

## Abbreviations

ACSF: artificial cerebro-spinal fluid
AP: area postrema
Bic: bicuculline methiodide
CD: control diet
CNQX: cyano-7-nitroquinoxaline-2,3-dione
Cv: coefficient of variation
DL-AP5: DL-2-Amino-5-phosphonopentanoic acid
DMV: dorsal motor nucleus of the vagus
DVC: dorsal vagal complex
EPSC: excitatory postsynaptic current
GLP-1: glucagon-like peptide 1
HFD: high fat diet
IPSC: inhibitory postsynaptic current
MEA: multi-electrode array
NDS: normal donkey serum
NPY: neuropeptide Y
NTS: nucleus of the solitary tract
OXA: orexin A
PBS: phosphate-buffered saline
PFA: paraformaldehyde
SCN: suprachiasmatic nuclei of the hypothalamus
SD: standard deviation
TTX: tetrodotoxin
ZT: Zeitgeber time

## ACKNOWLEDGEMENTS

Authors would like to thank Patrycjusz Nowik for excellent animal care.

## AUTHORS CONTRIBUTIONS

LC, KPC, JSJL, MHL and HDP conceived the study. LC and MHL supervised the study and provided financial support. LC and JDK designed, performed, analysed and interpreted electrophysiological studies with the help from JSJL. KP wrote custom MatLab and Spike2 scripts for automated spike- sorting and further analysis of multi-electrode array data. JSJL and JDK and performed spike-sorting. JDK and AMS performed immunohistochemical studies. JDK performed epifluorescence imaging. LC, HDP and JDK wrote the manuscript and all authors agreed to its final form.

## AVAILABILITY OF DATA AND MATERIALS

The data that support the findings of this study are available from the corresponding author upon reasonable request.

## SUPPLEMENTARY TABLES

**Table 1.**
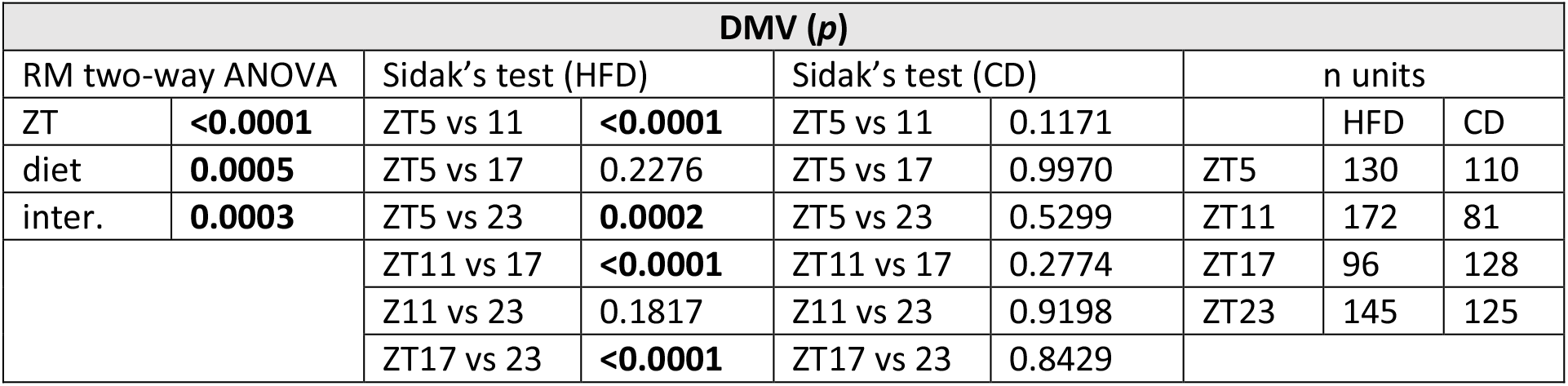
Statistical analysis for short-term recordings of spontaneous neuronal activity in the dorsal motor nucleus of the vagus (DMV) with multi-electrode arrays. CD – control diet, HFD – high-fat diet.

**Table 2.** Statistical analysis for synaptic activity in the dorsal motor nucleus of the vagus (DMV) in rats fed high-fat (HFD) control diet (CD). PSC –postsynaptic currents, total; EPSC – excitatory postsynaptic currents; IPSC – inhibitory postsynaptic currents

| Frequency |  |  |  |  |  |  |  |
| --- | --- | --- | --- | --- | --- | --- | --- |
| PSC |  |  |  | EPSC |  |  |  |
| two-way ANOVA (*p*) |  | n cells |  | two-way ANOVA (*p*) |  | n cells |  |
| ZT | **0.0443** | CD ZT6-12 | 14 | ZT | **0.0006** | CD ZT6-12 | 17 |
| diet | **0.0137** | CD ZT18-0 | 14 | diet | 0.0974 | CD ZT18-0 | 14 |
| inter. | 0.9150 | HFD ZT6-12 | 14 | inter. | 0.4937 | HFD ZT6-12 | 16 |
|  |  | HFD ZT18-0 | 12 |  |  | HFD ZT18-0 | 15 |
| IPSC |  |  |  |  |  |  |  |
| two-way ANOVA (*p*) |  | n cells |  | Sidak's test ZT6-12 – ZT18-0 (*p*) |  |  |  |
| ZT | **0.0018** | CD ZT6-12 | 13 | CD | 0.6321 |  |  |
| diet | **<0.0001** | CD ZT18-0 | 15 | HFD | **0.0005** |  |  |
| inter. | **0.0472** | HFD ZT6-12 | 16 |  |  |  |  |
|  |  | HFD ZT18-0 | 16 |  |  |  |  |
| Rise time |  |  |  |  |  |  |  |
| EPSC |  |  |  | IPSC |  |  |  |
| two-way ANOVA (*p*) |  | n cells |  | two-way ANOVA (*p*) |  | n cells |  |
| ZT | 0.0783 | CD ZT6-12 | 17 | ZT | **0.0035** | CD ZT6-12 | 13 |
| diet | 0.2759 | CD ZT18-0 | 14 | diet | 0.3498 | CD ZT18-0 | 15 |
| inter. | 0.0654 | HFD ZT6-12 | 16 | inter. | 0.5830 | HFD ZT6-12 | 16 |
|  |  | HFD ZT18-0 | 15 |  |  | HFD ZT18-0 | 16 |
| Decay time constant |  |  |  |  |  |  |  |
| EPSC |  |  |  | IPSC |  |  |  |
| two-way ANOVA (*p*) |  | n cells |  | two-way ANOVA (*p*) |  | n cells |  |
| ZT | 0.3050 | CD ZT6-12 | 17 | ZT | 0.4746 | CD ZT6-12 | 13 |
| diet | 0.0803 | CD ZT18-0 | 14 | diet | 0.3897 | CD ZT18-0 | 15 |
| inter. | 0.0692 | HFD ZT6-12 | 16 | inter. | 0.3055 | HFD ZT6-12 | 16 |
|  |  | HFD ZT18-0 | 15 |  |  | HFD ZT18-0 | 16 |

**Table 3.**
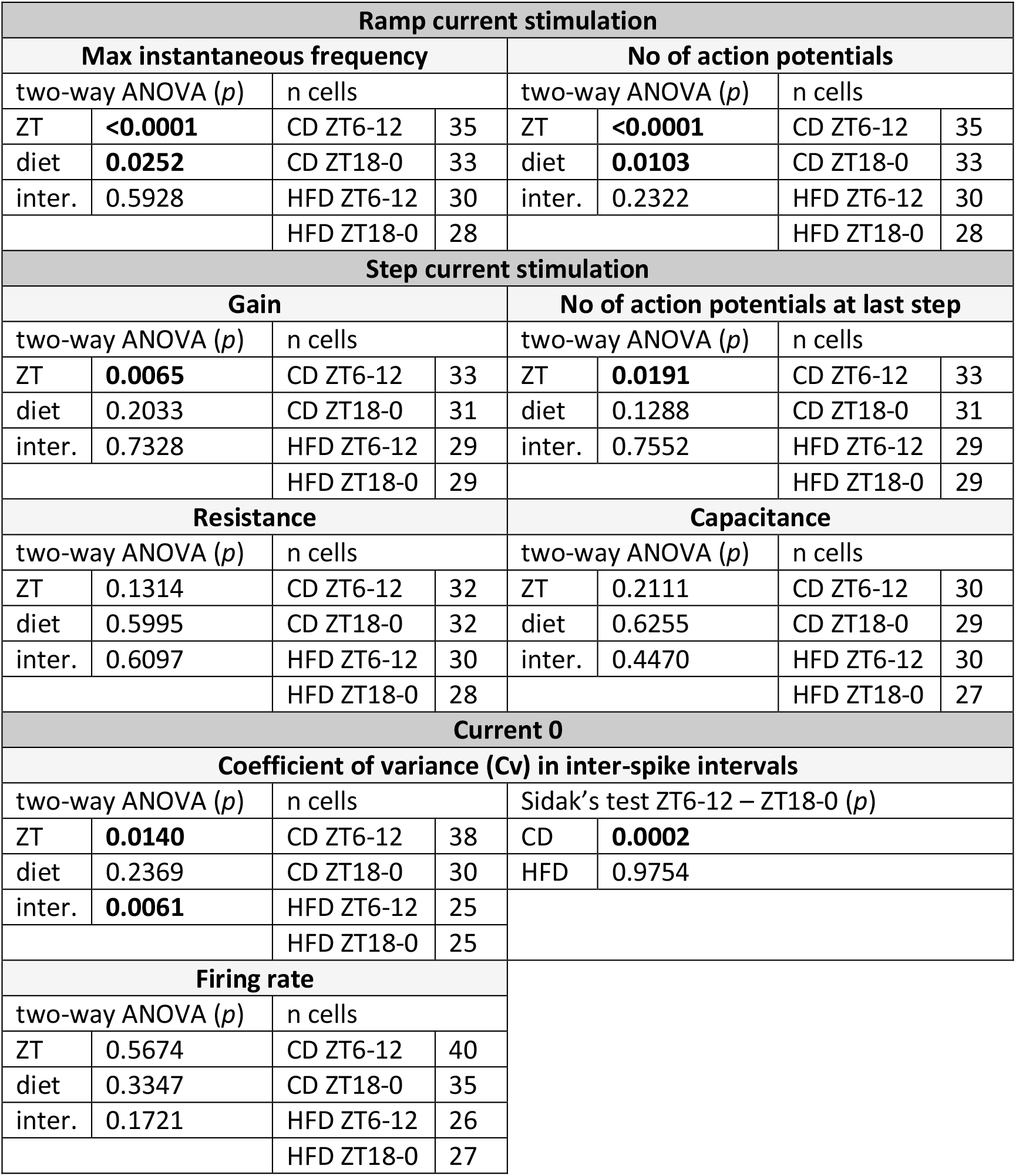
Statistical analysis for excitability tests in the dorsal motor nucleus of the vagus (DMV) in rats fed high-fat (HFD) control diet (CD).

**Table 4.** Statistical analysis for short-term recordings of the response amplitude to orexin A (OXA) and glucagon-like peptide 1 (GLP-1) in the dorsal vagal complex with multi-electrode arrays. CD – control diet, DMV – dorsal motor nucleus of the vagus, HFD – high-fat diet, NTS – nucleus of the solitary tract.

| OXA DMV (*p*) |  |  |  |  | GLP-1 DMV (*p*) |  |  |  |  |
| --- | --- | --- | --- | --- | --- | --- | --- | --- | --- |
| RM two-way ANOVA |  | n activated / inhibited / total |  |  | RM two-way ANOVA |  | n activated / inhibited / total |  |  |
| ZT | 0.3337 |  | HFD | CD | ZT | 0.8256 |  | HFD | CD |
| diet | 0.8543 | ZT5 | 105/3 /152 | 85/2 /124 | diet | 0.1012 | ZT5 | 75/2 /152 | 90/3 /124 |
| inter. | 0.3013 | ZT11 | 119/3 /178 | 34/0 /55 | inter. | 0.1498 | ZT11 | 106/4 /178 | 29/0 /55 |
|  |  | ZT17 | 72/3 /115 | 125 /1/162 |  |  | ZT17 | 64/3 /115 | 92/3 /162 |
|  |  | ZT23 | 92/6 /150 | 67/2 /109 |  |  | ZT23 | 74/1 /150 | 69/0 /109 |

**Table 5.** Statistical analysis for drug-evoked inward current amplitudes of the dorsal motor nucleus of the vagus (DMV) in rats fed high-fat (HFD) control diet (CD).

| OXA |  |  |  |  |  |  |  |
| --- | --- | --- | --- | --- | --- | --- | --- |
| -50 mV |  |  |  | -65 mV |  |  |  |
| two-way ANOVA (*p*) |  | n cells |  | two-way ANOVA (*p*) |  | n cells |  |
| ZT | **0.0015** | CD ZT6-12 | 14 | ZT | 0.6383 | CD ZT6-12 | 20 |
| diet | 0.9842 | CD ZT18-0 | 13 | diet | **0.0176** | CD ZT18-0 | 14 |
| inter. | 0.1215 | HFD ZT6-12 | 13 | inter. | 0.3810 | HFD ZT6-12 | 16 |
|  |  | HFD ZT18-0 | 13 |  |  | HFD ZT18-0 | 10 |
| GLP-1 |  |  |  |  |  |  |  |
| -50 mV |  |  |  |  |  |  |  |
| two-way ANOVA (*p*) |  | n cells |  | Sidak's test ZT6-12 – ZT18-0 (*p*) |  |  |  |
| ZT | 0.1410 | CD ZT6-12 | 9 | CD | **0.0272** |  |  |
| diet | 0.2441 | CD ZT18-0 | 9 | HFD | 0.8323 |  |  |
| inter. | **0.0306** | HFD ZT6-12 | 9 |  |  |  |  |
|  |  | HFD ZT18-0 | 11 |  |  |  |  |
| -65 mV |  |  |  |  |  |  |  |
| two-way ANOVA (*p*) |  | n cells |  |  |  |  |  |
| ZT | 0.7957 | CD ZT6-12 | 9 |  |  |  |  |
| diet | 0.8955 | CD ZT18-0 | 11 |  |  |  |  |
| inter. | 0.3779 | HFD ZT6-12 | 9 |  |  |  |  |
|  |  | HFD ZT18-0 | 8 |  |  |  |  |

